# Fluorescence-activated cell sorting of *Escherichia coli* L-forms reveals maintained synchronized nucleic acid synthesis dynamics

**DOI:** 10.64898/2026.09.21.753196

**Authors:** Marjolein E. Crooijmans, Johannes H. de Winde, Dennis Claessen

## Abstract

L-form bacteria are wall-deficient variants that proliferate without an intact cell wall and independently of the canonical cell division machinery. Despite increasing insights into how L-forms survive and proliferate without a cell wall, fundamental questions regarding how cell cycle processes, including DNA and RNA synthesis, are coordinated in the wall-less state remain poorly understood. Here, we demonstrate that fluorescence-activated cell sorting (FACS) provides a robust approach for quantitative analysis of *Escherichia coli* L-forms. Flow cytometry revealed two reversible subpopulations differing in intracellular nucleic acid content, which rapidly re-established their heterogeneous distribution following sorting. Remarkably, cells with initially low nucleic acid content underwent a synchronized, population-wide increase in nucleic acid synthesis within 24 hours, a phenomenon that was independently confirmed by time-lapse fluorescence microscopy. Analysis of liquid cultures stained with SYTO and DAPI dyes showed that this transient increase occurred during early exponential growth and was driven predominantly by RNA rather than DNA synthesis. Consistent with observations in walled *E. coli*, RNA levels peaked before declining as cultures transitioned towards nutrient limitation. Together, these findings establish FACS as a powerful tool for studying L-form biology and reveal that, despite the absence of an intact cell wall and canonical cell division, L-forms retain coordinated, population-wide regulation of nucleic acid synthesis.

**Importance:** Wall-deficient, or L-form, bacteria are increasingly recognized as biologically and medically relevant. L-forms can survive without the cell wall structure targeted by many commonly used antibiotics, and have been associated with persistent and recurrent infections, including recurrent urinary tract infections caused by *Escherichia coli* (Mickiewicz et al., 2019). Understanding how these unusual bacterial forms survive, grow and organize essential cellular processes is important for both fundamental microbiology and for understanding their potential clinical relevance. However, studying these cells is challenging because of their fragile nature. Reliable technologies that can analyze individual cells and separate distinct subpopulation are therefore needed. In this study, we investigate wall-deficient forms of *Escherichia coli* and demonstrate the usefulness of flow cytometry as a tool to study these cells. Expanding the methods available for studying wall-deficient bacteria will help clarify their biology and improve our understanding of their role in infection and treatment failure.

## Introduction

Bacterial proliferation requires tight coordination of chromosome replication and segregation with cell growth and division (Merrikh et al., 2011, Lang and Merrikh, 2018, Pountain et al., 2024, Olivi et al., 2025). In *Escherichia coli*, replication initiation is regulated by DnaA, while chromosome organization and division-site regulatory systems coordinate FtsZ-ring formation with chromosome segregation and promote its correct positioning at mid-cell. Cell elongation and septation are coupled to peptidoglycan (PG) synthesis through the activities of the elongasome and divisome, thereby linking the envelope biogenesis to the cell-cycle progression (Adams and Errington, 2009, Kleckner et al., 2018, Booth and Lewis, 2019). Together, these regulatory mechanisms ensure concerted genome inheritance and maintenance of cell morphology (Dewachter et al., 2018).

The *E. coli* cell envelope consists of an inner membrane, a PG layer in the periplasm and an asymmetric outer membrane containing lipopolysaccharides. PG provides mechanical strength and maintains cell shape. When PG synthesis is inhibited, cells become wall-deficient and generally lose the ability to proliferate (Tabata et al., 2019). Under suitable conditions, however, wall-deficient cells can proliferate as L-forms. Unlike walled cells, L-forms divide independently of FtsZ through membrane deformation and scissions, potentially resembling proposed reproductive mechanisms of primitive early life forms (Errington et al., 2016, Studer et al., 2016, Mercier et al., 2016, Errington, 2017, Kanaparthi et al., 2021, Lubbers and Claessen, 2025).

How other cellular processes are organized during L-form proliferation remains poorly understood. In particular, it is unclear whether nucleic acid synthesis retains temporal organization when cells lack a cell wall and no longer divide through regulated binary fission. Addressing this question requires methods that can quantify nucleic acid content and population heterogeneity at the single-cell level.

Fluorescence-activated cell sorting (FACS) offers a powerful approach to address these questions. As a flow cytometry-based technique, FACS enables measuring and sorting of individual cells according to their physical characteristics, such as size and shape, and with fluorescent markers reporting cellular properties including nucleic acid content (Herzenberg et al., 2002, Dangl and Lanier, 2013). However, application of FACS to L-forms presents several technical challenges. The absence of a protective cell wall renders cells highly fragile and dependent on osmoprotective conditions for survival (Osawa and Erickson, 2019, Mercier et al., 2014). During FACS analysis, cells are suspended in a sheath fluid that is typically based on phosphate-buffered saline (PBS), which may be insufficiently osmoprotective for wall-deficient cells. Furthermore, the sorting process exposes the cells to substantial pressure and shear forces that can reduce post-sorting viability (Pfister et al., 2020).

Here, we established a FACS approach for analyzing and sorting *E. coli* L-forms and used it to examine nucleic acid synthesis during growth. Despite the absence of PG-mediated growth and FtsZ-dependent division, L-form populations showed reproducible growth-phase-associated changes in nucleic acid content, consistent with observations in walled cells, indicating that organization of cellular activity persists during L-form proliferation.

## Results

### Establishing FACS as a platform to study L-form cell-cycle dynamics

To investigate whether nucleic acid synthesis remains coordinated in wall-less *E. coli*, we first established a FACS-based workflow suitable for fragile L-form cells. Because the absence of a cell wall makes L-forms highly susceptible to osmotic and mechanical stress, it was critical to determine whether these cells could survive the sorting procedure while retaining viability for downstream analyses.

Figure 1A presents three strains of *E. coli*, an L-form (LTE0), its evolved strain (LTE800) and its parental strain (wild type) which can grow in a walled (rod or Rev) and wall-less (spheroplasts or L-form) state depending on the growth media (Crooijmans et al., 2025) (Table 1). Morphology was verified using microscopy of overnight cultures of each of these conditions, revealing rod-shaped cells in LB and spherical cells in LPB supplemented with penicillin G (penG) (Fig. 1B). To assess whether wall-deficient bacteria, including spheroplasts and L-forms, remained viable after cell sorting, we sorted the lowest and highest 25% of cells based on forward scatter (FSC) from each overnight culture. Subsequently, these sorted samples were plated on LB agar and LMPA supplemented with penG to test for their recovery and viability post-sorting (Fig. 1C). All samples were able to recover on LB agar. On LPMA supplemented with penG, neither walled nor wall-deficient wild-type cells (Rod^WT^ and Spheroplast^WT^) survived the sorting procedure, as no viable colonies were recovered. In contrast, both evolved L-form lineages (LTE0 and LTE800) showed partial viability after sorting, with colony-forming efficiencies ranging from 0.001% for L-form^LTE0^ to 18% for Rev^LTE800^ (Fig. 1D). As expected, the walled (Rev^LTE800^) and wall-less (L-form^LTE800^) cells of the evolved LTE800 strain had a higher recovery rate compared to the LTE0 lineage (Crooijmans et al., 2025).

**Figure 1.**
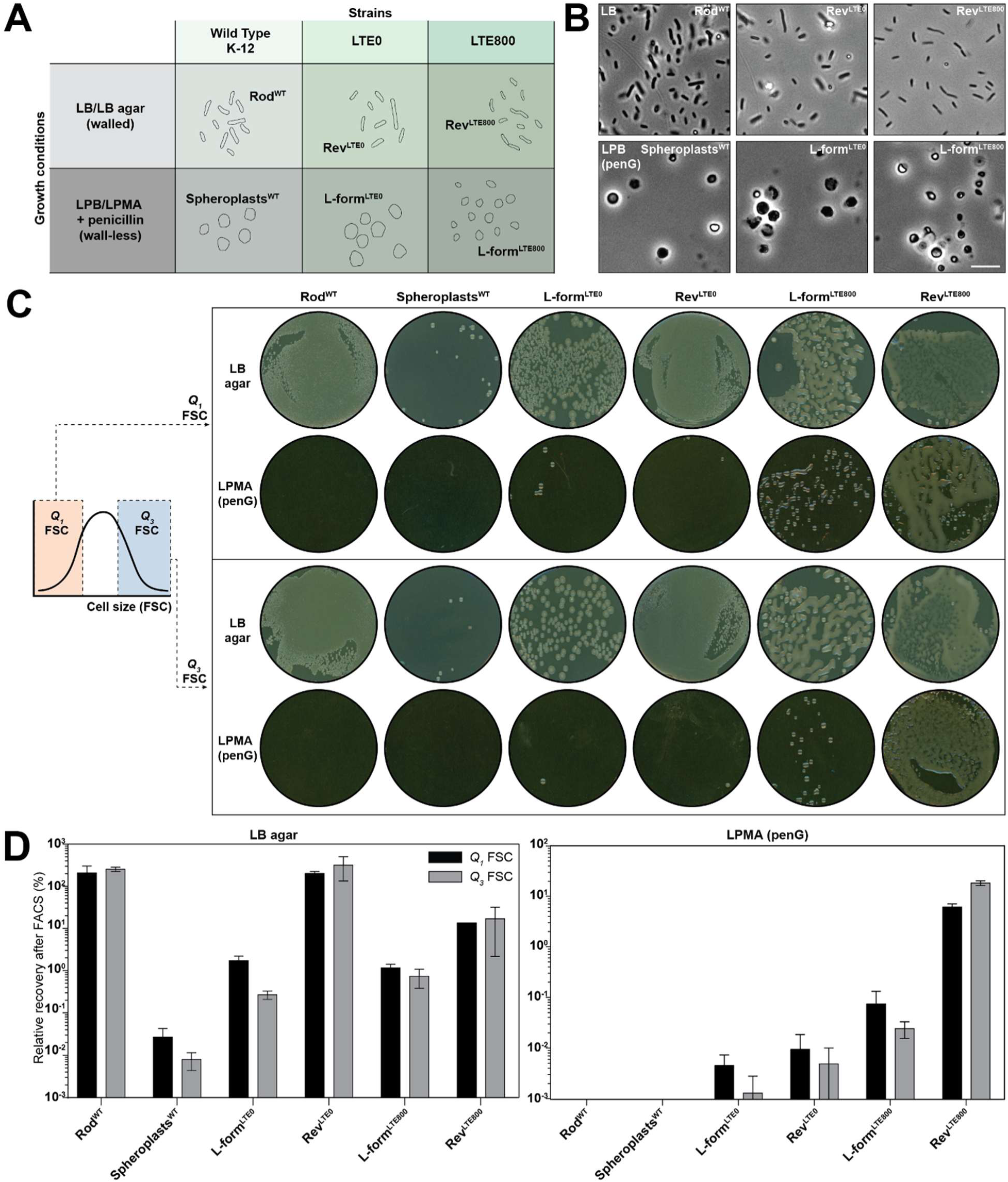
Wall-less cells remain viable after cell sorting. **A)** Overview of three *E. coli* strains under two conditions. For walled growth, cells were grown in LB or LB agar and are called Rods for WT and revertants (Rev) for the LTE lineages. Wall-less growth as spheroplasts for WT and L-forms for LTE was achieved by growing cells in osmoprotective medium (LPB medium for liquid cultures and LPMA medium for growth on agar plates) supplemented with penicillin G. More details on the strains can be found in Table 1. **B)** Wild-type, Rev^LTE0^ and Rev^LTE800^ cells were cultured in LB medium, while spheroplasts, L-form^LTE0^ and L-form^LTE800^ were grown in LPB medium. Overnight cultures were imaged using transmission light microscopy. Scale bar is 10µm. **C)** Cell populations were sorted based on lowest 25% (Q_1_) and highest 25 (Q_3_) FSC, 100.000 sorting events were plated on LB agar and LPMA (penG) media and imaged after 48 hours incubation at 37°C. **D)** Quantification of viable cells following sorting was determined by colony counts on plates and presented as bar plots.

**Table 1.** Strains of *E. coli* used in this study.

| Lineage | Genotype | Walled state (LB) | Wall-less state (LPB/LPMA) | Source |
| --- | --- | --- | --- | --- |
| Wild Type | K-12 MG1655. F<sup>-</sup>. λ<sup>-</sup>. ilvG<sup>-</sup> rfb-50. rph-1. pUA139-serW::gfpmut2 | Rod<sup>WT</sup> | Spheroplasts<sup>WT</sup> | CGSC 6300 (Zaslaver et al., 2006) |
| LTE0 | L-form mutant of wild type. pUA139-intF::gfpmut2 | Rev<sup>LTE0</sup> | L-form<sup>LTE0</sup> | (Crooijmans et al., 2025) |
| LTE800 | Long-term evolution mutant of L-form/Rev<sup>LTE0</sup>. pUA139-intF::gfpmut2 | Rev<sup>LTE800</sup> | L-form<sup>LTE800</sup> | (Crooijmans et al., 2025) |

### Analysis of nucleic acid content reveals two distinct L-form populations

To investigate nucleic acid dynamics in wall-less *E. coli*, we focused on the evolved L-form strain LTE800, which displayed the highest post-sorting viability and was therefore most suitable for downstream analyses. Cells were stained with the fluorescent nucleic acid dye SYTO 85 and analyzed by flow cytometry. The FSC versus side scatter (SSC) plot of L-form^LTE800^ cells showed that most cells clustered within a similar size and internal complexity/density range (Supplemental Fig. 1A). Density plots of SYTO fluorescence, however, revealed a heterogeneous distribution of nucleic acid signal, separating the culture into a major population with low fluorescence and a smaller population with high fluorescence (Fig. 2A). This suggested the presence of distinct nucleic acid states within the L-form population. Because larger cells are expected to contain more nucleic acids, we next asked whether the two SYTO populations simply reflected differences in cell size. Cells were computationally gated into the lowest (Q1) and highest (Q3) quartiles of SYTO fluorescence and compared for FSC, as a proxy for cell size. The Q3 population exhibited significantly higher FSC values than Q1 (median FSC 103,000 versus 39,000 A.U.; Wilcoxon test, *p* < 2.2 × 10⁻¹⁶), indicating that cells with higher SYTO fluorescence are generally larger (Fig. 2B). Furthermore, the FSC distribution of Q3 cells was considerably broader, exhibiting a 151-fold greater variance than that of Q1 cells (Fig. 2C), demonstrating that the high-SYTO population is substantially more heterogeneous in size.

**Figure 2.**
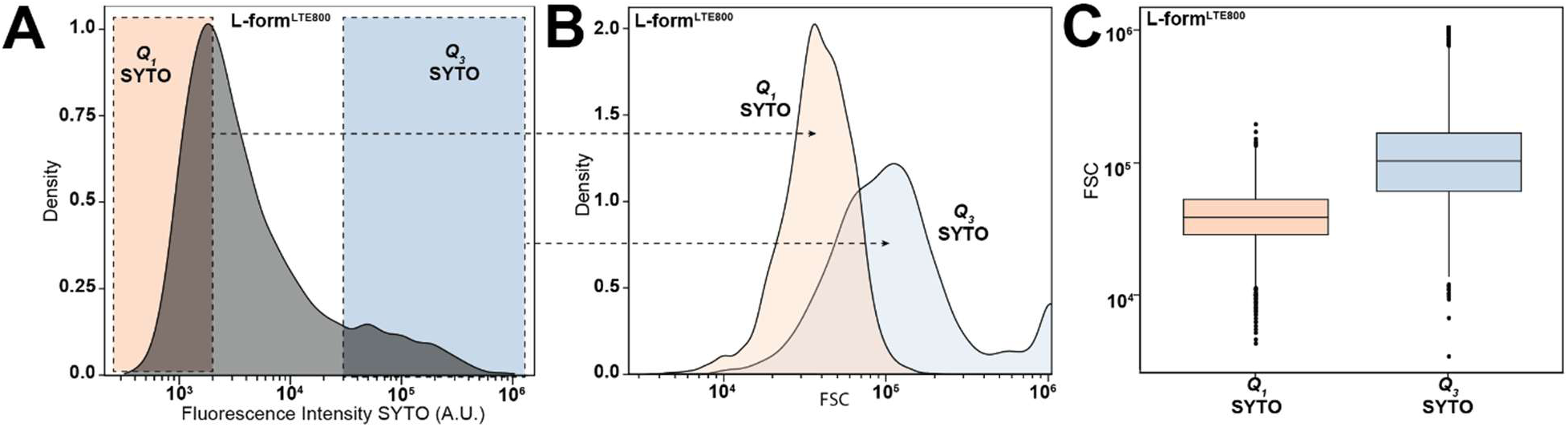
Flow cytometry analysis of L-form^LTE800^ cells revealed heterogeneous nucleic acid distribution. **A)** Density plot illustrating the distribution of SYTO fluorescence intensities. Based on SYTO fluorescent intensities, L-form^LTE800^ cells in the bottom and top quantiles (25%) were computationally classified into two groups: Q_1_ SYTO and Q_3_ SYTO, respectively. **B)** Density plot comparing the FSC of the cells in the Q_1_ and Q_3_ categories. **C)** Box and whisker plot showing median, interquartile range and whiskers representing the FSC spread of cells in the Q_1_ and Q_3_ categories. The whiskers represent data ranging within the 1.5 interquartile range values. Outliers outside this range are plotted as individual dots. Q_1_ median is 38,668, n = 2,501, Q_3_ median is 103,001, n = 2,500.

To determine whether the observed differences in SYTO fluorescence could be explained solely by cell size, we examined the relationship between SYTO fluorescence and FSC. Raw SYTO intensity showed a moderate positive correlation with FSC (Pearson’s *r* = 0.54; Supplemental Fig. 1B), confirming that larger cells generally contain more nucleic acids. After regression-based correction for cell size, however, this relationship was almost completely abolished (*r* = 0.07; Supplemental Fig. 1C). Together, these analyses show that although larger L-form cells generally contain more nucleic acids, cell size alone does not account for the observed heterogeneity in SYTO fluorescence. The presence of two SYTO populations therefore reflects differences in relative nucleic acid content rather than simply differences in cell size.

### Nucleic acid-defined subpopulations re-establish population heterogeneity after sorting

To determine whether the low- and high-SYTO populations represent stable subpopulations or dynamic physiological states, L-form^LTE800^ cells were sorted into the lowest (Q1) and highest (Q3) quartiles of SYTO fluorescence. Sorting accuracy was confirmed by re-analysis of a subset of sorted cells. The Q1 and Q3 populations were subsequently cultured in LPB medium (Fig. 3A). The low-SYTO (Q1) population responded rapidly after sorting. Within 24 hours, the median SYTO fluorescence increased 4.4-fold (from 3,752 to 16,445 A.U.), indicating a population-wide increase in nucleic acid synthesis. After 48 hours, fluorescence heterogeneity began to reappear and, by 72 hours (OD600 ≈ 1.5), the characteristic bimodal distribution was fully re-established.

**Figure 3.**
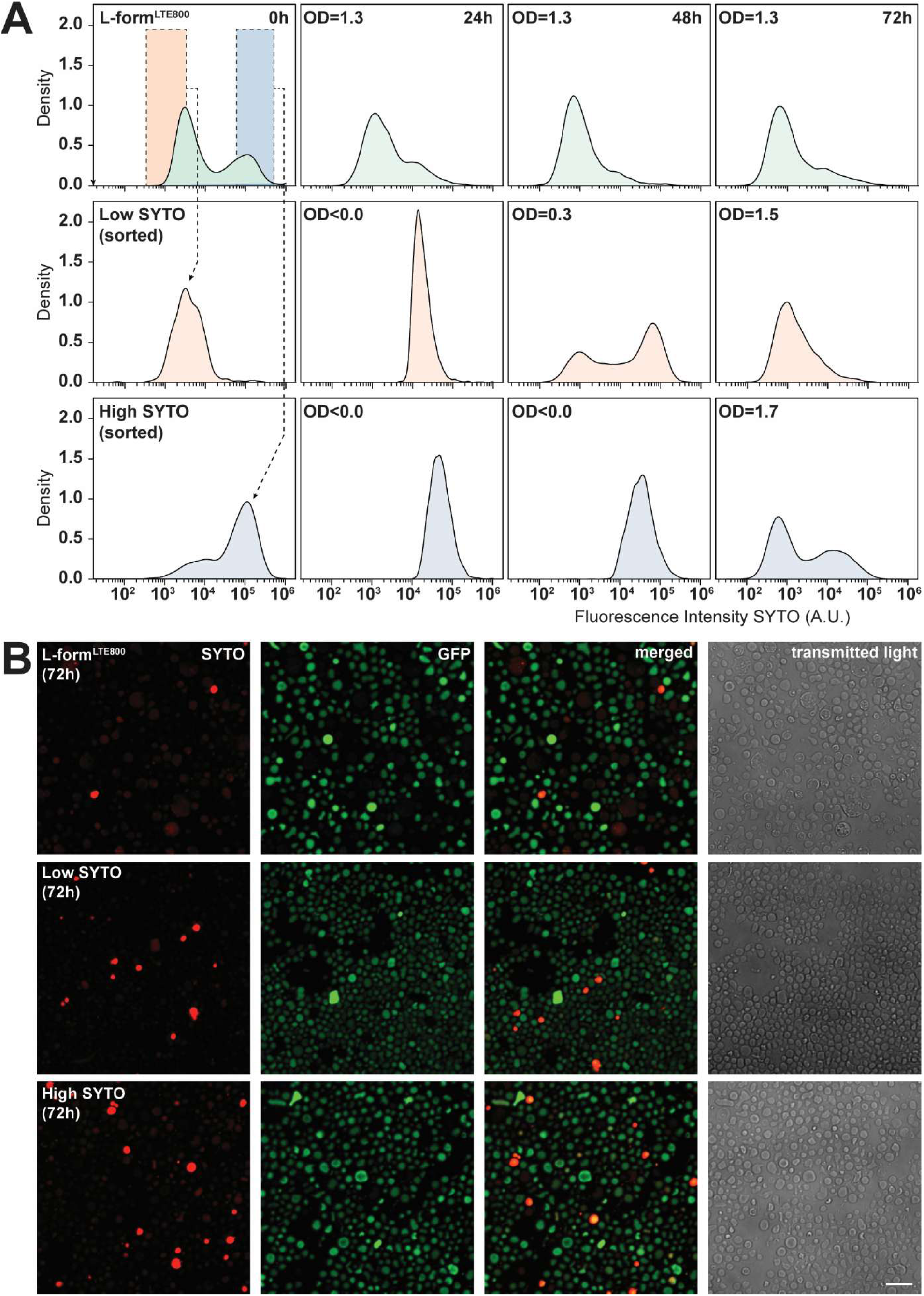
Dynamics of L-form^LTE800^ cells sorted by nucleic acid content over time. **A)** Density plots of flow cytometry data showing L-form^LTE800^ cells sorted into low (Q_1_) and high (Q_3_) SYTO populations using FACS and cultured in liquid medium. Measurements were taken at 0, 24, 48 and 72 hours. Non-sorted L-form^LTE800^ cells were included as a control. Fluorescence intensities are in logarithmic scale. Data in arbitrary units. **B)** Fluorescence confocal images of L-form^LTE800^ cells Q_1_ and Q_3_ populations after 72 hours of growth. Scale bar is 10µm.

In contrast, the high SYTO (Q_3_) population showed little growth during the first 72 hours after sorting, as reflected by the low OD_600_ measurements. During this period, cells retained their high SYTO fluorescence. Once growth resumed, however, the original heterogeneous distribution likewise re-emerged, resulting in the same two-population profile observed in the unsorted culture. Fluorescence microscopy performed after 72 hours confirmed that the unsorted, Q1 and Q3 cultures were morphologically similar and all displayed comparable heterogeneous nucleic acid distributions (Fig. 3B). Together, these findings demonstrate that the low- and high-SYTO populations are not genetically fixed subpopulations but reversible physiological states. Regardless of their initial nucleic acid content, both sorted populations regenerated the characteristic heterogeneous distribution, indicating that nucleic acid levels are dynamically regulated at the population level.

### Time-lapse imaging reveals colony-wide synchronization of nucleic acid synthesis

The reversible transition between low- and high-SYTO states suggested that nucleic acid synthesis remains dynamically regulated in wall-less *E. coli*. To determine whether this transition occurs in a coordinated manner across the population, we monitored SYTO fluorescence in growing L-form^LTE800^ colonies by time-lapse fluorescence microscopy. The initial microscopy images showed a heterogeneous population consisting predominantly of cells with low SYTO fluorescence interspersed with a smaller number of highly fluorescent cells, mirroring the distribution observed by flow cytometry (Fig. 4A). After approximately 2 hours of growth, however, the entire colony displayed a simultaneous increase in SYTO fluorescence. This coordinated shift suggests that nucleic acid synthesis is synchronously activated across the wall-deficient population, consistent with the rapid increase in SYTO signal observed during regrowth of the sorted Q1 population (Fig. 4A, Fig. 3A). Quantification of the mean fluorescence intensity showed that colony-wide SYTO fluorescence reached a maximum after approximately 3 hours of growth. Subsequently, fluorescence gradually declined while the colony underwent a burst of daughter cell release, suggesting that the transient increase in nucleic acid synthesis precedes active proliferation (Fig. 4B). Interestingly, cells that initially displayed high SYTO fluorescence retained this elevated signal after 4 hours, whereas fluorescence in the remaining cells gradually declined. Combined with the flow cytometry and cell-sorting experiments, these results reveal that wall-deficient *E. coli* L-forms transiently synchronize nucleic acid synthesis at the population level before returning to heterogeneous nucleic acid states during continued growth. This coordinated behavior demonstrates that loss of the cell wall does not abolish temporal regulation of nucleic acid synthesis, but instead that wall-less cells retain an organized growth-stage-dependent program.

**Figure 4.**
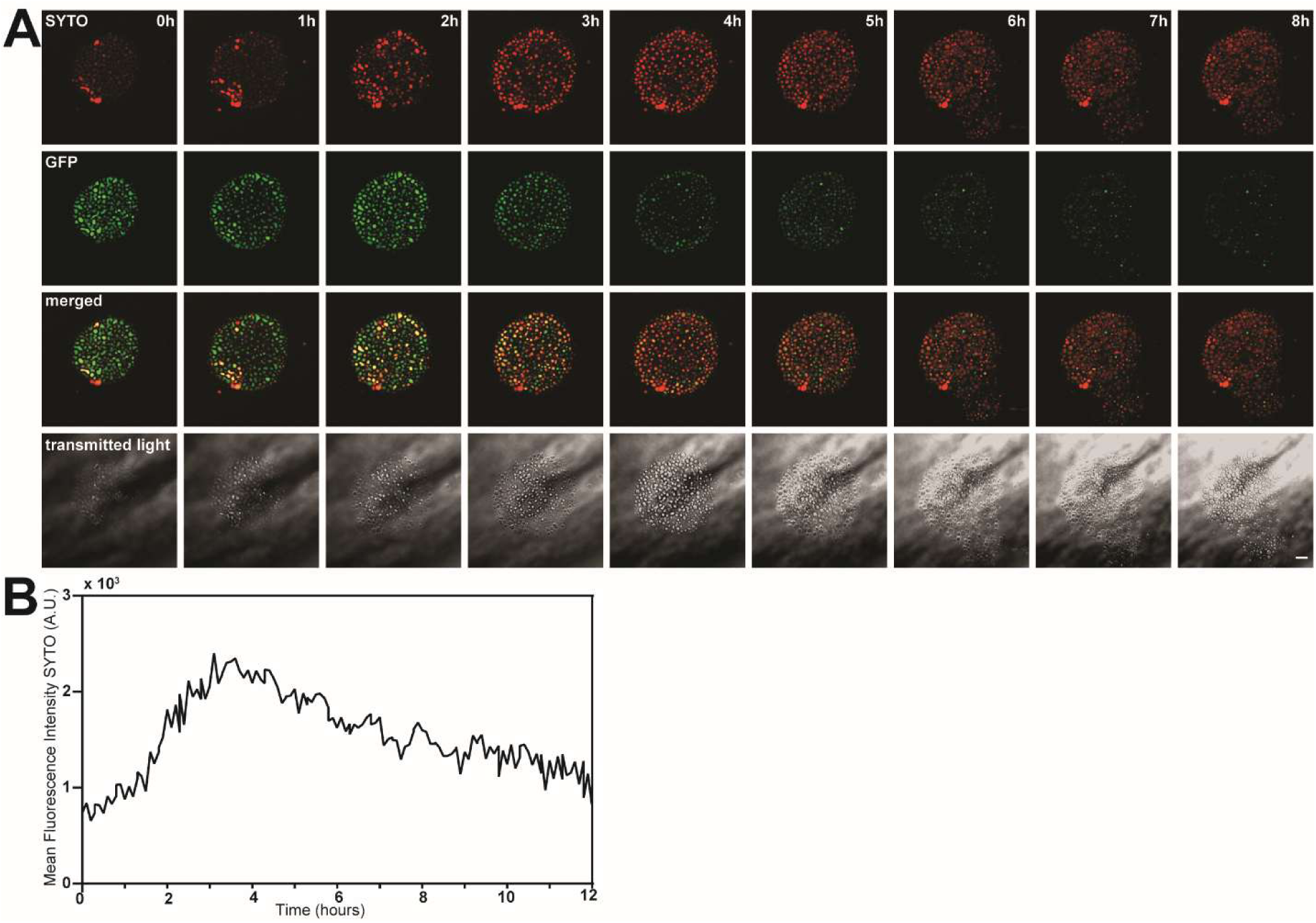
Colony-wide nucleic acid-associated fluorescence varies across the growth of L-form^LTE800^ cells. **A)** Time-lapse confocal images of a L-form^LTE800^ colony grown under an LPMA pad supplemented with SYTO dye, acquired at hourly intervals (Supplementary Video 1). Scale bar is 10µm. **B)** Quantification of mean fluorescence intensity of the L-form^LTE800^ colony measured every 5 minutes, expressed in arbitrary units.

### Synchronized nucleic acid synthesis accelerates during early exponential growth

To determine whether the synchronized increase in nucleic acid synthesis is linked to the growth stage of wall-less *E. coli*, L-form^LTE800^ was cultured in liquid medium and sampled every 1.5 hours throughout growth (Fig. 5A). To avoid photobleaching and RNA degradation, each sample was stained and imaged immediately after harvesting; specifically with SYTO to detect total nucleic acids, and with DAPI to specifically label double-stranded DNA. During the earliest growth stages (OD600 ≈ 0.03–0.07), the population consisted predominantly of cells with low SYTO fluorescence, similar to the initial distributions observed by flow cytometry and time-lapse microscopy. As cultures entered early exponential growth (OD600 ≈ 0.1–0.3), SYTO fluorescence increased sharply across the population before gradually declining at higher cell densities, recapitulating the transient burst of nucleic acid synthesis observed in the previous experiments (Fig. 5A-B).

**Figure 5.**
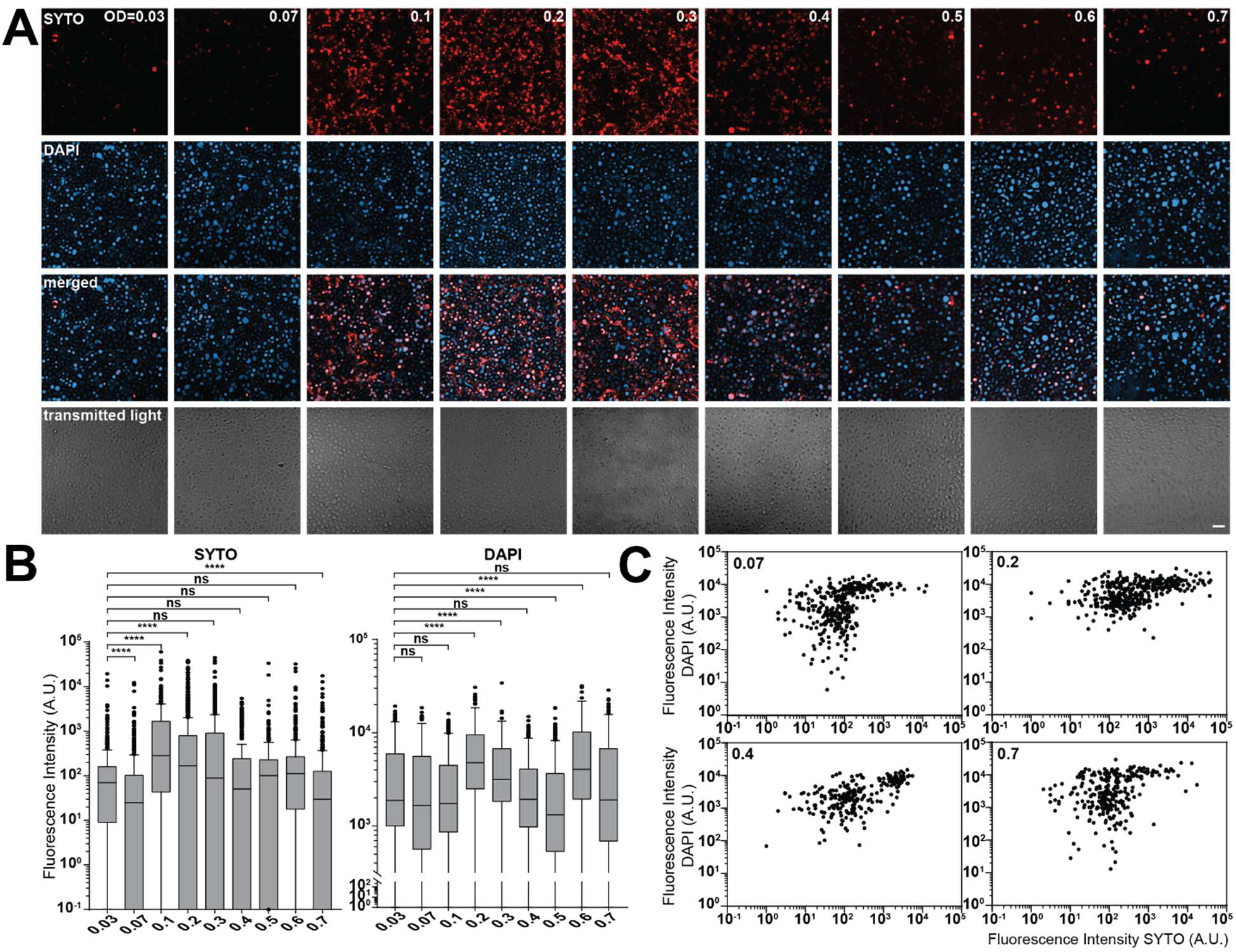
SYTO fluorescence is increased in early exponential growth of L-form^LTE800^ cells. **A)** Fluorescence confocal images of L-form^LTE800^ cultures at various optical densities (OD_600_). Scale bar is 10µm. **B)** Box and whisker plot showing the median, interquartile range and whiskers representing the SYTO and DAPI fluorescence intensities after background corrections per cell at each OD_600_ measurement. Whiskers represent the data ranging within the 1.5 interquartile range values. Outliers outside this range are plotted as individual dots. Fluorescence intensities are in arbitrary units. **C)** Scatter plots of single-cell SYTO and DAPI fluorescence intensities across four OD_600_ conditions.

In contrast, DAPI fluorescence remained relatively constant throughout growth, with only modest increases observed around OD600 = 0.1 and 0.6 (Fig. 5B). Quantification of individual-cell fluorescence confirmed that changes in SYTO and DAPI signals followed distinct temporal patterns. Between OD600 = 0.07 and 0.1, mean SYTO fluorescence increased 11.4-fold, whereas DAPI fluorescence increased only marginally by 1.05-fold. Subsequently, DAPI fluorescence increased more strongly during later growth intervals (OD600 = 0.1–0.2: 2.7-fold versus 0.6-fold for SYTO; OD600 = 0.5–0.6: 3.1-fold versus 1.1-fold for SYTO) (Fig. 5B). As expected, an identical colony-wide increase in nucleic acid synthesis (SYTO) at OD = 0.2-0.3 was also observed in wild type *E. coli* cells grown in LB medium (Supplemental Fig. 2).

These observations indicate that the synchronized burst in SYTO fluorescence during early exponential growth cannot be explained solely by DNA replication. Instead, the majority of the transient increase likely reflects increased RNA accumulation, while DNA synthesis follows a distinct temporal pattern. To further examine the relationship between DNA content and total nucleic acid content, we compared DAPI and SYTO fluorescence at different growth stages (Fig. 5C). Spearman’s rank correlation revealed only weak associations during early (OD600 = 0.07; ρs = 0.384) and late (OD600 = 0.6; ρs = 0.306) growth, whereas a substantially stronger correlation was observed during early exponential growth (OD600 = 0.1; ρs = 0.598; *p* < 0.0001 for all comparisons). This transient increase in correlation suggests that the heterogeneous nucleic acid states identified earlier become temporarily synchronized as the population enters exponential growth before diverging again during later growth. Together, these findings demonstrate that wall-less *E. coli* L-forms undergo a coordinated, growth-stage-dependent burst of nucleic acid synthesis that is driven predominantly by RNA accumulation. Although DNA synthesis also increases during growth, its temporal dynamics differ from those of total nucleic acid synthesis, indicating that RNA production is the principal contributor to the synchronized increase in SYTO fluorescence.

## Discussion

In this study, we demonstrate that wall-deficient *E. coli* cells can be successfully sorted using FACS and reveal substantial heterogeneity in nucleic acid content within stable L-form^LTE800^ populations. Nucleic acid content increased population-wide during early exponential growth and subsequently declined. This change appears to be predominantly associated with RNA rather than DNA synthesis. Our findings raise three main points for discussion: the coordination of nucleic acid synthesis in wall-deficient cells, the physiological basis and reversibility of nucleic acid heterogeneity, and the opportunities and limitations of FACS for studying L-form biology.

Despite the loss of the cell wall and disruption of cell division, L-form cells retained coordinated nucleic acid synthesis, suggesting that key aspects of the cell cycle remain functional. We found a colony-wide increase in nucleic acid content during early exponential growth, followed by a decrease during later growth (Fig. 4A, Fig. 5A). The fact that this pattern was observed across the population and that sorted cells were able to re-establish the same population-wide heterogeneity after isolation, suggests that nucleic acid synthesis is not a consequence of random differences between individual cells, but rather coupled to the growth state of the culture (Fig. 3A). Similar observations were seen in walled *E. coli* cells (Supplemental Fig. 2). In walled cells, RNA synthesis is closely linked to nutrient availability and growth rate. During growth in LB, growth rate and cell mass decrease from approximately OD600= 0.3 as nutrients become limiting (Sezonov et al., 2007). This triggers the ppGpp-mediated starvation response, which leads to an immediate inhibition of almost all RNA synthesis (Jin et al., 2012). Rapid RNA degradation has also been observed during stationary and starvation phases (Gummesson et al., 2020, Sorensen et al., 2018). These growth- and nutrient-dependent regulatory processes that shape nucleic acid synthesis in walled *E. coli* may similarly account for the in- and decrease in RNA content and re-emergence of a heterogeneous SYTO distribution observed in L-form^LTE800^ cells. However, as these studies were performed in LB medium, it remains unclear how the nutrient composition of LPB contributes to this response.

The heterogeneous nucleic acid content may reflect the existence of two populations of cells that are physiologically different within the culture. Using flow cytometry, we identified a population of high and low SYTO-fluorescence (Fig. 2). This heterogeneity was restored within 72h after the respective populations were sorted and grown (Fig. 3). This suggests cells can transition between these physiological states. Interestingly, optical density measurements showed that cells with high SYTO fluorescence recovered more slowly after sorting than low-SYTO populations. One possible explanation is that the high-SYTO population represents cells undergoing active RNA synthesis, which may leave them temporarily more sensitive to the physical stress associated with sorting. Nevertheless, the re-establishment of the heterogeneous population after sorting suggests that nucleic acid distribution is a dynamic feature in the growing L-form population. It remains to be established if other cell cycle-related processes are also coordinated in cells lacking their cell wall.

Besides providing insights into L-form physiology, our results demonstrate for the first time that wall-deficient *E. coli* cells can be sorted in the L-form state, despite their increased fragility (Fig. 1C-D). This provides opportunities to investigate L-form heterogeneity by separating cells according to physical traits such as size and shape, but also physiological characters as nucleic acid, DNA and membrane content. The sorted cells can be subsequently followed to study their growth, morphology, gene expression or other phenotypes. However, the low survival rate of wall-deficient cells introduces a potential recovery bias, as cells that better withstand the mechanical and fluidic stresses of sorting are more likely to survive. This should be considered when interpreting sorted populations, particularly as cells from the high-SYTO fraction required a longer recovery period after sorting (Fig. 3A). Importantly, the heterogeneous SYTO distribution re-emerged during subsequent growth, suggesting that this pattern reflects a reversible state of cellular activity rather than a persistent sorting bias. Nevertheless, the ability to sort viable L-form cells substantially expands the experimental toolbox for studying wall-deficient bacteria.

## Methods

### Growth media

*E. coli* cells were grown in two types of media to control cell wall formation. Luria Bertani (LB and LB agar) broth was used for the growth of walled cells, while the osmostable L-form medium (LPB for liquid, LPMA for agar) was used for growth of cells without their cell wall (Ramijan et al., 2018). All cells were grown at 37°C, and media were supplemented with kanamycin (50 µl ml^-1^). OD_600_ measurements were performed using an Ultrospec™ 2100 pro UV-Visible spectrophotometer.

### Bacterial strains

This study uses three *E. coli* lineages (Fig. 1A, Table 1). All strains carry a GFP marker and kanamycin resistance cassette (Crooijmans et al., 2025).

### Flow cytometry and Fluorescent Assisted Cell Sorting (FACS)

Quantification of FSC, SSC and SYTO stains were performed using a Bio-Rad S3e Fluorescence-activated Cell Sorter with ProSort Software, version 1.6. ProLine Universal Calibration Beads (Bio-Rad Laboratories) were used to calibrate the machine. As shealth fluid, ProFlow Sort Grade 8x Sheath Fluid was used.

#### Sample preparations

Cells of Rod^WT^, Spheroplasts^WT^, Rev^LTE0^, L-form^LTE0^, Rev^LTE800^ and L-form^LTE800^ were grown overnight in LB or LPB medium, respectively. To remove debris, 1 ml of cells were spun down at 3,200 rcf for 2 min (Eppendorf 5424), washed twice in P-buffer (Kieser et al., 2000) and filtered using a pluriStrainer (40 µm). If required, 20 µl of filtered cells were stained with 0.5 µl of SYTO 85 (5.0 mM, Invitrogen). Subsequently 20 µl of cells were diluted in 500 µl P-buffer and transferred to a 5ml sample tube (12/75 mm, round bottom, Greiner Bio-One).

#### Acquiring flow cytometry data

To be able to compare data across multiple time-points, the prepared cells were always measured using FSC (voltage 266), SSC (voltage 225), GFP (voltage 747, 488 nm, 525/30 nm), SYTO (voltage 561, 561 nm, 586/25 nm). To minimize background noise, events were gated using the GFP signal (threshold 1.00). A minimum number of 10,000 events were measured.

#### Cell sorting

Prior to sorting, collection tubes (12/75 mm, round bottom, Greiner Bio-One) were pre-filled with 500 µl of P-buffer. Sorting gates were established to isolate the highest and lowest quantiles (25%) of the target population (see Fig. 1C). For FSC-based sorting, sorting was performed until 800,000 events were collected and all sorted cells were maintained on ice until further handling. To test post-sorting viability, a dilution series ranging from 100 to 100,000 sorted events was plated in triplicate onto vented 90-mm Petri dishes (Greiner BIO-ONE) containing LB and LPMA (penG). Plates were incubated at 37°C for 48 hours. Viability was then quantified by counting individual colony-forming units on each plate. Additionally, the plates inoculated with 100,000 events were imaged using an Epson Perfection V600 scanner (Fig. 1C). For the SYTO-based sorting, a total of 400,000 cells were sorted. To verify sorting purity, an aliquot of 1,000 cells was re-analyzed (Fig. 3A). The remaining cells were transferred to 20 ml LPB medium and incubated at 37°C.

#### Analysis and visualization

Flow cytometry standard (.fcs) files were imported and analyzed in Rstudio (version 2025.9.2, R language (R-Core-Team, 2018) using the *flowCore* package (Hahne et al., 2009). Raw data from each sample was extracted and concentrated into one dataset using *dplyr* for comparative analysis (Wickham et al., 2023). For data visualization, scatter plots and density plots were generated using *ggplot2* (Wickham, 2016). All parameters were log10-transformed. For scatter plots comparing FSC (FSC-AREA), SSC (SSC-AREA), GFP (FL1-AREA) and SYTO (FL2-AREA), color scaled to indicate event density using the *viridis* packages (Garnier et al., 2024). To calculate the descriptive statistics of each group, the n(), mean(), sd(), min(), quantile(), median(), max() were performed in R. OpenAI’s ChatGPT GPT-5.5 was used to troubleshoot R code.

### Microscopy

#### Transmitted light microscopy

To image the morphological differences between the six strains, cells were grown overnight, after which 5 µl of the cultures were transferred to a glass microscopy slide and covered with a glass cover slip. Images were taken with a Zeiss Axio Lab A1 A-plan 40x/0.65 Ph2 microscope equipped with a Zeiss Axiocam 105 camera and Zeiss 2.5 Lite software.

#### Fluorescence microscopy

Confocal microscopy was performed using an inverted Zeiss LSM 900 confocal microscope equipped with an Airyscan 2 module and a C-Apochromat 63x/1.20 W Korr UV-VIS-IR objective. Images acquisition was controlled using Zeiss Zen 3.1 software (Blue Edition). For single-timepoint imaging, sample preparation was adjusted based on optical density to ensure similar cell counts per image. For high-density cultures (OD600 >0.5), 500 µl of culture was centrifuged at 3200 rcf and supernatant was removed leaving a final volume of 20 µl. For low-density cultures (OD600 <0.5), up to 10 ml of culture was pelleted at 3,200 rcf and similarly concentrated to 20 µl. When required, samples were fluorescently stained by adding 0.5 µl of SYTO 85 (5.0 mM, Invitrogen) and 0.5 µl of DAPI (14.3 mM, Invitrogen), followed by a 15 min incubation at room temperature. The cells were spotted onto an LMPA pad, incubated for an additional 15 min and flipped upside down on an 8-well μ-Slide (Ibidi). For colony time-lapse microscopy, sorted cells from the lowest 25% SYTO-stained L-form^LTE800^ population were prepared as described above. The slide was incubated at 37°C for 24 h to allow the sorted cells to recover. To maintain a constant fluorescent signal over time, the agar pad was supplemented with a 0.1x concentration of SYTO dye. A viable colony was then imaged every 5 minutes for 12 h with a z-sections of 0.10 µm, maintained at 37°C. The same laser excitation and emission settings were used across all experiments DAPI (353 nm, 465 nm), GFP (488 nm, 509 nm) and SYTO (578 nm, 598 nm).

#### Analysis and image preparation

Following acquisition, images were stored in OMERO and prepared for publication using OMERO.figure (Allan et al., 2012). All quantitative image analysis was performed using Fiji (Schindelin et al., 2012). To measure the mean SYTO fluorescence intensity of the whole colony during the time-lapse imaging, channels were first split to isolate the SYTO signal. Maximum intensity projections were generated from the z-stacks, after which the colony region was selected using the oval selection tool to measure the intensity across the entire time-lapse. For the single cell analysis, the center of each bacterial cell was marked on the transmitted light images using the multi point tool to extract the corresponding SYTO and DAPI fluorescence intensities. A minimum of 500 cells were analyzed per condition. To correct background noise, the fluorescence of 50 intercellular regions was measured for each condition and subtracted of the raw cellular values. Statistical analysis and data visualization (bar and scatter plots) were performed in Prism. To evaluate the relationship between SYTO and DAPI signals, a Spearman rank correlation was used, as this non-parametric test is more robust for single cell fluorescence data. For cell size diameters, transmission light microscopy images were imported to Fiji, threshold was set to remove background noise, colors were inverted and using *Analyze Particles*, *Fit ellipse* was used to get the *Major* score, the major axis length. The data was plotted using Graphpad Prism (version 10.0). For multiple comparisons, the non-parametric Krustal-Wallis test was used, followed by Dunn’s multiple comparison test. Ns = P > 0.05, * = P ≤ 0.05, ** = P ≤ 0.01, *** = P ≤ 0.001, **** = P ≤ 0.0001.

## Data availability

Source data, including raw measurements, analysis scripts, and processed data, are available from the corresponding author upon request.

## Acknowledgements

We thank Bio-Rad for technical support and troubleshooting of the FACS machine. We are also grateful to Joost Willemse for suggesting the use of and lending us the SYTO 85 dye, which largely improved the experimental workflow. This work was funded by a VICI grant (no. VI.C.192.002) from the Dutch Research Council to Dennis Claessen. M.C. designed the experimental setup, performed the FACS and confocal microscopy experiments, analyzed the data and wrote the original draft of the manuscript. M.C., J.H.W., and D.C. contributed to the editorial process.

